# CRISPRi-based circuits for genetic computation in plants

**DOI:** 10.1101/2022.07.01.498372

**Authors:** Muhammad Adil Khan, Gabrielle Herring, Marina Oliva, Elliott Fourie, Jia Yuan Zhu, Benjamin Johnston, Jahnvi Pflüger, Tessa Swain, Christian Pflüger, James Lloyd, David Secco, Ian Small, Brendan Kidd, Ryan Lister

**Affiliations:** Australian Research Council Centre of Excellence in Plant Energy Biology, School of Molecular Sciences, The University of Western Australia, WA, Australia; Harry Perkins Institute of Medical Research, The University of Western Australia, WA, Australia; CSIRO Synthetic Biology Future Science Platform, QLD, Australia

## Abstract

Synthetic gene circuits can enable new cellular behaviours by integrating multiple input signals into customisable genetic programs. However, gene circuit development in plants has been limited by a lack of orthogonal and modular parts required for their construction. Here, we present a tool-kit of reversible CRISPRi-based gene circuits for use in plants. First, we created a range of engineered repressible promoters of different strengths and used them as integrators for the construction of NOT and NOR gates in *Arabidopsis* cells. Next, we determined the optimal processing system to express sgRNAs from RNA Pol II promoters to introduce NOR gate programmability and interface it with host regulatory sequences. Finally, we connected multiple NOR gates together in layered arrangements to create OR, NIMPLY, and AND logic functions. Our CRISPRi circuits are orthogonal, compact, reversible, programmable, and modular, providing a new platform for sophisticated and deliberate spatio-temporal control of gene expression in plants.

## Introduction

To engineer plants for improved stress tolerance or growth and yield traits, plant biotechnology relies heavily on the expression of introduced transgenes to produce desired phenotypes and cellular activities. However, the expression of transgenes with strong constitutive promoters can lead to silencing, metabolic burden, or other deleterious effects on yield and, as a result, the benefit from the desired transgene may not be fully realised^1–5^. To further improve the toolset for controlling plant gene expression, it would be valuable to produce customizable expression programs that not only act within a certain spatiotemporal condition, but also integrate modifiers to alter the phenotype depending on the overall condition of the plant. By incorporating multiple signalling cues, a sophisticated gene expression program could, for example, either boost an endogenous stress response or alternatively manage yield-associated traits according to the “threat level” of the stress^6–9^.

Synthetic biology offers the potential to address these challenges through the use of synthetic gene circuits. Analogous to biological gene regulatory networks, synthetic gene circuits aim to integrate multiple input signals into a core logic function to control the output promoter activity in an input state-dependent manner, thereby adding more sophisticated capabilities for constructing genetic programs and controlling expression^6,7,10–12^. Synthetic gene circuits are able to integrate multiple signals to produce Boolean logic operations such as NOR, OR, and AND to ultimately produce an output in the form of a change in transcription in the presence of the desired combination of inputs. Building a set of robust and orthogonal Boolean logic operations would allow for logic gates to be stacked together. This would enable the development of more complex circuits and provide far greater capabilities to fine tune plant pathways or enhance cell functions based on user-defined inputs.

To build gene circuits, recombinases and transcriptional regulators obtained from other host organisms are often used, however refactored proteins may be limited in diversity or may not function as expected in plants, and thus require performance and cross-reactivity testing for each intended plant species. Nevertheless, gene circuits have been constructed using recombinase proteins or orthogonal transcription factors (TFs) in plants^13–18^. Alternatively, plant TF domains can be combined with orthogonal DNA binding domains and synthetic promoters to create logic^13,19^, however the use of plant TF domains requires endogenous regulatory factors, and therefore are difficult to program beyond their natural range of activity.

As an alternative to using recombinases and TFs, circuits based on CRISPR interference (CRISPRi) that use a nuclease-dead Cas9 (dCas9) protein to inhibit target promoter activity have been successfully implemented in *Escherichia coli, Saccharomyces cerevisiae*, and mammalian cells for the development of synthetic gene circuits^20–26^. In plants, CRISPRi has been demonstrated to work for transcriptional regulation^27–30^, but to date it has not been leveraged to construct synthetic gene circuits. Circuit construction using CRISPRi offers multiple potential advantages, including having dynamically reversible activity states, being easily expandable (by adding more sgRNA constructs to build layered logic gates), and having the ability to simultaneously target multiple endogenous loci without modification or the need to add recognition elements in promoters. Therefore, to take advantage of these features, we have developed a toolkit for dCas9-based logic gate construction in plants, and in the process outlined key design requirements for effective regulation of dCas9-based circuits in plants.

## Results

### Designing an integrator for implementing NOR logic in plants

To construct a plant gene circuit, we first aimed to build a NOR gate, which is a functionally complete (“universal”) logic gate, meaning that it can be combined with other NOR gates to produce all sixteen 1- or 2-input Boolean logic gate functions. The inputs and outputs of a circuit can be represented in a truth table using values of 1 (ON) and 0 (OFF). A NOR gate maintains an ON state in the absence of an input signal, but is switched to the OFF state when either input A or B is present (Fig. 1a). Therefore, in the presence of a single input (A or B) a NOR gate will act as a NOT gate. To create a dCas9-based NOR gate in plants, two different sgRNAs (input A and input B) are used to recruit dCas9 to bind to corresponding target sites in an engineered promoter that drives the expression of the output gene. We refer to such an engineered promoter as an “integrator” for its ability to receive and integrate input signals (sgRNAs targeting dCas9) to control the corresponding output. Therefore, in the presence of input A, input B, or both inputs A and B, the dCas9 protein is targeted to the integrator and inhibits transcription initiation through CRISPRi, leading to repression of the output gene (Fig. 1a).

**Figure 1:**
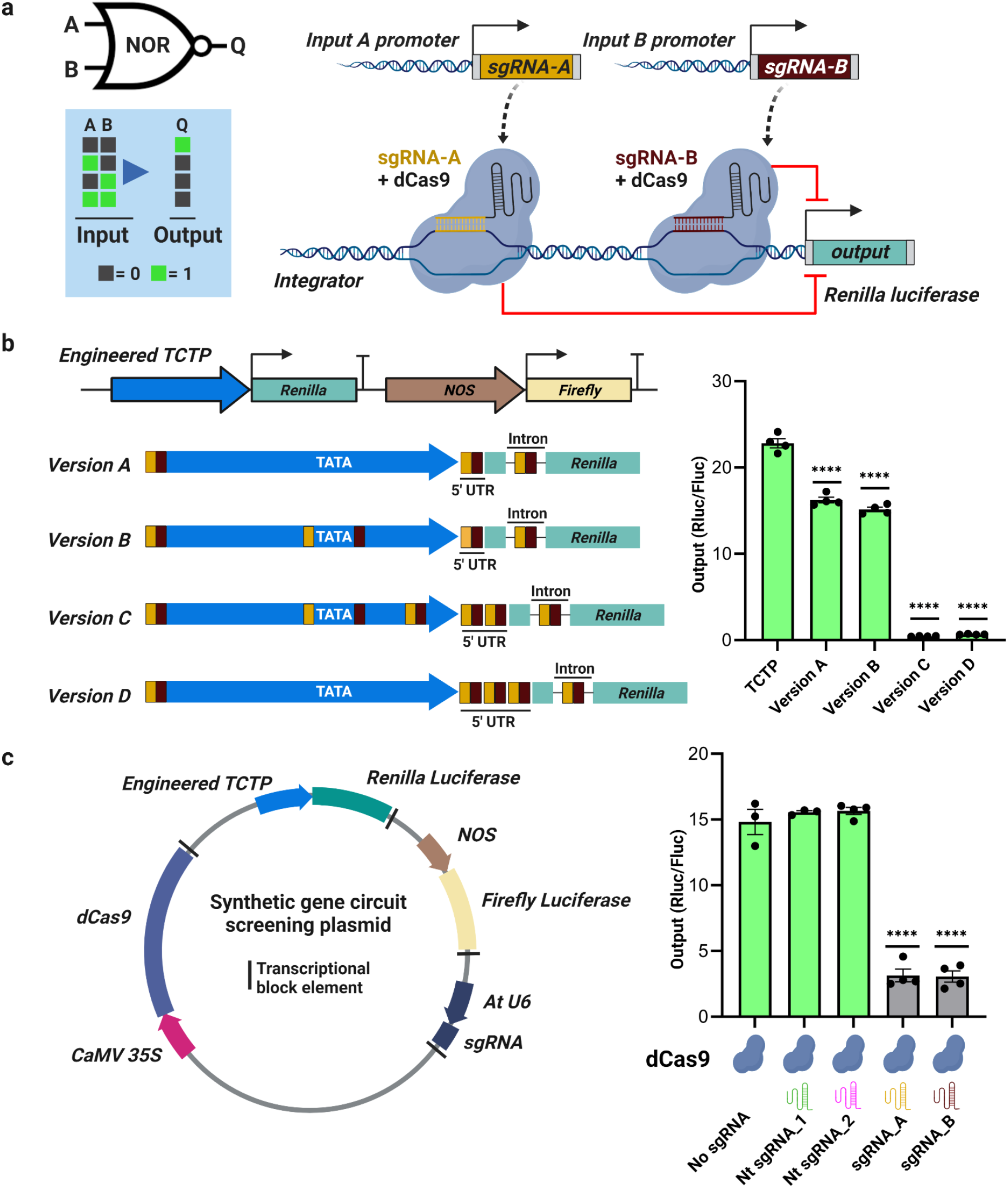
Designing and testing integrators for a plant CRISPRi/dCas9-based NOR gate. **(a)** Schematic representation, truth table for a 2-input NOR gate, and overview of a dCas9-based NOR logic gate. In the presence of sgRNA-A, sgRNA-B, or both sgRNAs, the dCas9 protein binds to the integrator and by CRISPRi represses transcription of *Renilla* luciferase, which is the logic gate output. **(b)** Schematic of the construct(s) used for testing the activity of engineered *TCTP* promoters driving the expression of *Renilla* luciferase, where the *NOS* promoter driving the expression of firefly luciferase was used as a transfection control. Different versions of the engineered *TCTP* promoter are shown in blue, with yellow and brown boxes representing the positions of sgRNA-A and sgRNA-B binding sites, respectively (not to scale), and green boxes representing the *Renilla* luciferase coding sequence. Activities of Versions A-D of the engineered *TCTP* promoter in *Arabidopsis* protoplasts 16 hours post transfection (hpt). Promoter activity is measured as the ratio of *Renilla* luciferase derived luminescence (Rluc) to firefly luciferase derived luminescence (Fluc), and termed output (Rluc/Fluc, y-axis). Error bars = standard error, n = 4. **(c)** Schematic representation of a synthetic gene circuit screening plasmid containing all the components (different versions contained different sgRNA sequence or lacked sgRNA sequence). Effect of input sgRNA-A or sgRNA-B on the integrator activity compared to a No sgRNA and Non-targeted (Nt) control sgRNAs, all in the presence of dCas9, using Version B of the engineered *TCTP* promoter. Output (relative luciferase activity [Rluc/Fluc]) was measured 24 hpt, and shown with the same y-axis used in panel B. Error bars = standard error, n = 3-4. Asterisks (**** = p ≤ 0.0001) indicate a significant difference using a one-way ANOVA and post-hoc test (Tukey multiple comparison test).

To enable rapid testing of logic gates in plants, we developed a high-throughput dual luciferase (*Renilla*:firefly) reporter assay to test circuit components in *Arabidopsis thaliana* leaf protoplasts. All circuit components were encoded on a single plasmid, eliminating the challenges of co-transfection. Developing a reliable high-throughput protoplast transfection assay was critical to increase testing throughput and reduce variation in protoplast transfection rates that would cause significant challenges in interpreting logic gate function^14^.

As a starting point to engineer a promoter for the NOR gate as an integrator, we selected the constitutively expressed *Arabidopsis TRANSLATIONALLY CONTROLLED TUMOR PROTEIN* (*TCTP*; 303 bp) promoter^31^, which has a moderate expression level relative to strong promoters such as the widely used cauliflower mosaic virus 35S mRNA (*CaMV 35S*) promoter^32^. Engineering of the promoter with unique synthetic sgRNA binding sites for dCas9 is required to avoid targeting the endogenous *TCTP* promoter and allows for the creation of diverse new promoters that can be repressed via CRISPRi. Therefore, we created four variants (Version A-D) of the *TCTP* promoter by inserting two distinct sgRNA binding site sequences, which replaced the native sequence of the same length, in different locations within the promoter, 5’ untranslated region (5’ UTR), as well as in the potato *IV2* intron^33^ inserted into the *Renilla reniformis* luciferase reporter gene (Fig. 1b). Version A had sgRNA binding sequences upstream of the promoter and in the 5’ UTR and intron, leaving the promoter free of sgRNA binding sites. Version B was the same as Version A except sgRNA binding sites were added immediately upstream and downstream of the TATA box. Version C and Version D had an additional sgRNA binding site inserted at the 3’ end of the *TCTP* promoter and an additional sgRNA binding site in the 5’UTR. Furthermore, Version D had an unmodified *TCTP* promoter sequence but had three sgRNA binding sites in the 5’UTR, one in the intron, and one sgRNA binding site immediately upstream of the promoter. To determine whether the introduction of sgRNA binding sites would affect the strength of the *TCTP* promoter in the absence of dCas9, each engineered *TCTP* promoter was fused to the *Renilla* luciferase (Rluc) coding sequence and cloned into a plasmid containing a firefly luciferase (Fluc) gene from *Photinis pyralis*, driven by the *NOPALINE SYNTHASE* (*NOS*) promoter, which functions as a normalizer for plasmid transfection. We then transfected these plasmids into protoplasts isolated from the leaves of *Arabidopsis* plants and performed dual luciferase assays to measure the promoter output, as inferred from the relative luminescence (Rluc/Fluc) detected. Versions A and B of the engineered *TCTP* promoter produced outputs (relative luminescence, Rluc/Fluc) that were 71% and 66% of the wild-type *TCTP* promoter, respectively, however the promoter activity was almost completely eliminated in Versions C and D (Fig. 1b). As the integrator must retain the ability to drive transcription in the absence of targeting sgRNAs, we chose Version B promoter to be tested for CRISPRi in the presence of sgRNAs.

To determine whether we could repress the Version B promoter by CRISPRi, we expressed dCas9 from the *CaMV 35S* promoter and used the *A. thaliana U6 SMALL NUCLEOLAR RNA26* (*AtU6-26*) promoter to express the input sgRNAs, all encoded in a single plasmid, followed by transfection into *Arabidopsis* leaf protoplasts (Fig. 1c). Both input sgRNA-A and input sgRNA-B effectively repressed the Version B promoter to ∼20% of the activity of the Version B promoter when No input sgRNA was present. The activity of the Version B promoter was similar in the absence of any sgRNA or when non-targeting (Nt) sgRNA-1 or -2 were included, which targeted the *Arabidopsis PDS* promoter or was a random sequence with no match in the *Arabidopsis* genome, respectively. Given the ability of this engineered promoter to be effectively repressed by input sgRNA signals, we refer to it as an integrator.

### NOR gate expansion and optimization

As general transcription factors and the RNA polymerase II holoenzyme complex bind to the promoter region surrounding the TATA box and transcriptional start site, this region should provide an effective loci for achieving maximum repression with CRISPRi, as demonstrated previously in other species^20–22^. Thus, we wanted to confirm whether all four copies of both sgRNA-A and sgRNA-B binding sites in the Version B integrator were required for effective CRISPRi. We therefore targeted sgRNA-A to different positions within the Version B integrator to determine the effect on CRISPRi. Our results show that CRISPRi is strongest when the sgRNA target sites are inserted immediately flanking the TATA box of the *TCTP* promoter, with one site 5’ and the other site 3’ of the TATA box, while targeting the 5’ UTR (+125 bp of TSS) was the next best location (Supplementary Fig. 1). Targeting the intron of *Renilla luciferase* (+282 bp) or immediately upstream of the integrator (−287 bp) failed to repress the reporter gene (Supplementary Fig. 1). Targeting the TATA box and the 5’ UTR at the same time had a slightly lower average repression level but was not statistically different from the TATA box only. As targeting the TATA box achieved similar levels of repression to the Version B integrator containing four pairs of the two sgRNA targeting sites, we chose to build subsequent logic gates with sgRNA target sites flanking the TATA box, as this simplified the construction of synthetic NOR gate integrators.

To expand the set of integrators we created two different libraries of engineered promoters based on the *TCTP* and *CaMV 35S* promoters by replacing the native sequences flanking the TATA box within the promoter with two distinct sgRNA binding sequences (Supplementary Tables 1 and 2; Fig. 2a). Interestingly, this generated a range of promoters with different strengths (Fig. 2b and c), demonstrating how variation of TATA box flanking sequences can cause significant variation in promoter strength. Therefore, this provides a viable approach to produce a wide range of integrators with different output strengths in plants.

**Figure 2:**
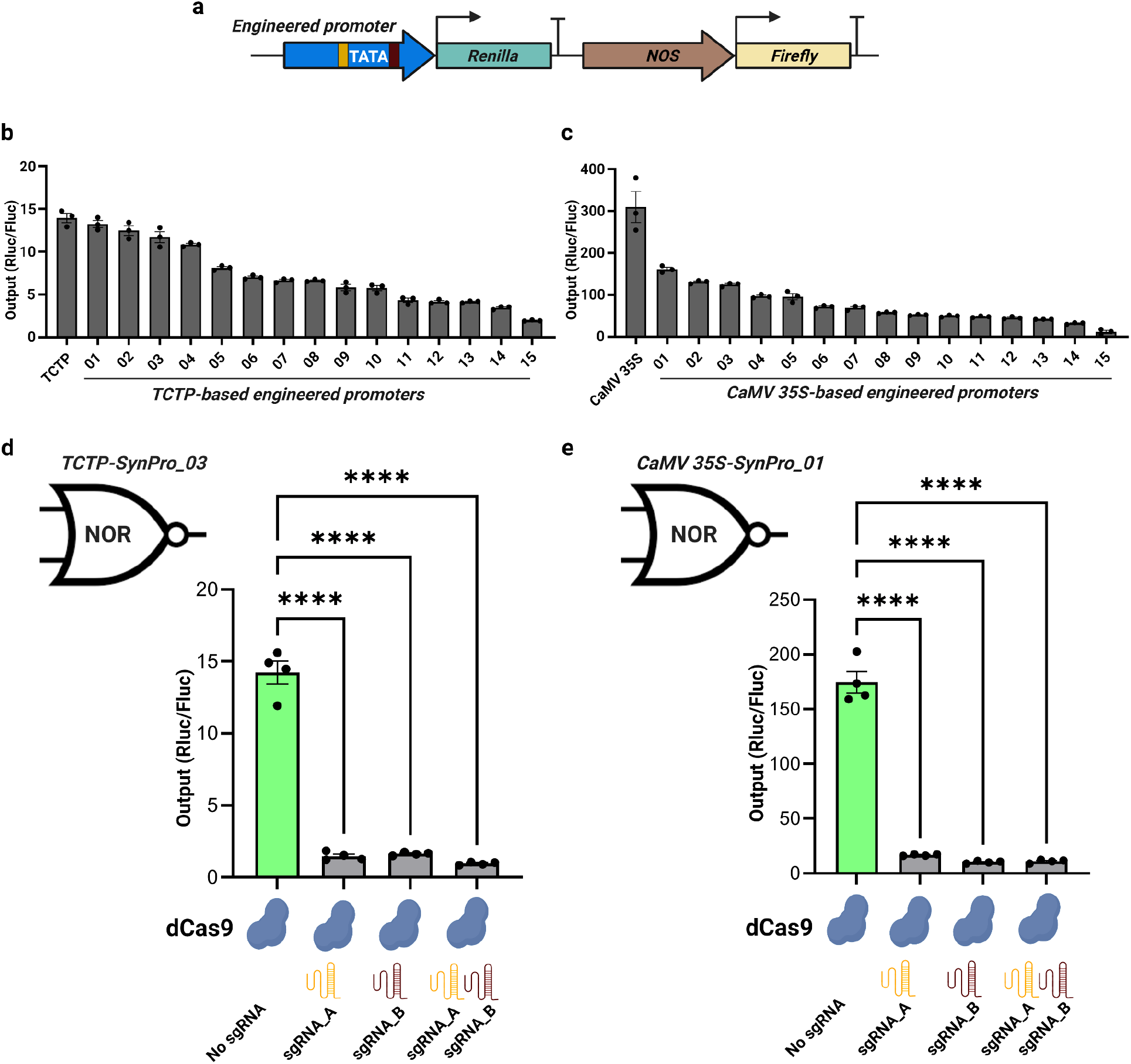
Testing libraries of *TCTP-* and *35S*-based synthetic promoters followed by the construction of NOR gates. **(a)** Schematic of the construct used for testing the activity of synthetic promoters in plant cells. The *TCTP*- and *CaMV35S*-based engineered promoters (yellow and brown boxes represent unique sgRNA binding sites) were used to drive the expression of *Renilla* luciferase (output), with firefly luciferase driven by the *NOS* promoter allowing normalization for variable transfection efficiency, and measurement of output (Rluc/Fluc). Activity of the library of synthetic **(b)** *TCTP*-based engineered promoters and **(c)** *CaMV 35S*-based engineered promoters in plant cells (n = 3). Circuit output for a CRISPRi-based functional NOR gate in plant cells using **(d)** *TCTP-SynPro_03* or **(e)** *CaMV 35S-SynPro_01* as the integrator, and different combinations of input sgRNAs, or No sgRNA. Error bars represent standard error (n = 4). Asterisks (**** = p ≤ 0.0001) indicate a significant difference compared to the No sgRNA using one-way ANOVA and post hoc test (Tukey multiple comparison test).

We next created new *TCTP*- and *CaMV 35S*-based NOR gates using integrators from these two different libraries. Different versions of the constructs that contained no sgRNA, sgRNA-A, sgRNA-B, or both sgRNAs -A and -B (Supplementary Fig. 2) were transfected into *Arabidopsis* protoplasts to test NOR gate activity at 24 hpt. We chose *TCTP-SynPro_03* and *CaMV 35S-SynPro_01* to act as integrators for these NOR gates because after sgRNA target site incorporation they showed activity relatively close to their unmodified native promoters (Fig. 2b and c). On average, we achieved up to 93% and 94% repression for the *TCTP-SynPro_03* and *CaMV 35S-SynPro_01* integrators, respectively, when either or both input sgRNAs were present compared to the No sgRNA control (Fig. 2d and e). Together, these findings demonstrate that the strength of logic gate integrators can be fine-tuned by altering TATA-flanking sgRNA target sequences, that CRISPRi is sufficient for repressing plant promoters of different strengths, and represent the first implementation of CRISPRi-based NOR gates in plants.

### Expanding NOR gate programmability

Ideally, the NOR gate should be programmable in response to endogenous or exogenous input signals to provide more sophisticated spatiotemporal control of gene expression *in planta*. Therefore, it is important to determine an effective sgRNA processing strategy for the expression of input sgRNAs from cell type-specific and inducible promoters, which typically utilise RNA Pol II, rather than the conventional RNA Pol III *U6* promoter. Unlike expression from a Pol III promoter, sgRNAs expressed from a Pol II promoter require additional processing to remove the 5’ cap and polyA tail. We therefore tested three different sgRNA processing systems: ribozymes, tRNAs, and Csy4. We tested a 5’ hammerhead ribozyme (HH) sequence and a 3’ Hepatitis Delta Virus ribozyme (HDV) sequence flanking the sgRNA sequence^34^ (Fig. 3a). We also tested the use of tRNA(Gly) sequences to flank the sgRNA sequence that are recognised by the endogenous RNase P and RNase Z to release a mature sgRNA^35,36,34^ (Fig. 3b). In contrast to the previous two approaches that are functional within a plant cell without additional factors, we also tested the CRISPR RNA endonuclease Csy4, which recognizes a specific 20 base sequence and cleaves the 3’ end of the recognition sequence. We introduced the Csy4-encoding gene as well as Csy4 recognition sequences upstream and downstream of the sgRNA sequence to express a mature sgRNA via a Pol II promoter^35,37,38^ (Fig. 3c). We cloned these sequences downstream of the *CaMV 35S* promoter as an example Pol II promoter, and compared the ability of ribozymes, tRNAs, and Csy4 to express the input sgRNAs targeting a *TCTP-SynPro_03*-based NOR gate. For each sgRNA processing system, different gene circuit construct versions were generated that contained no sgRNA, sgRNA-A, sgRNA-B, or both sgRNAs -A and -B, and were transfected into *Arabidopsis* protoplasts to test NOR gate activity 24 hpt. As with all other experiments, for each condition all circuit components were encoded on a single plasmid (similar to the constructs shown in Supplementary Fig. 2).

**Figure 3:**
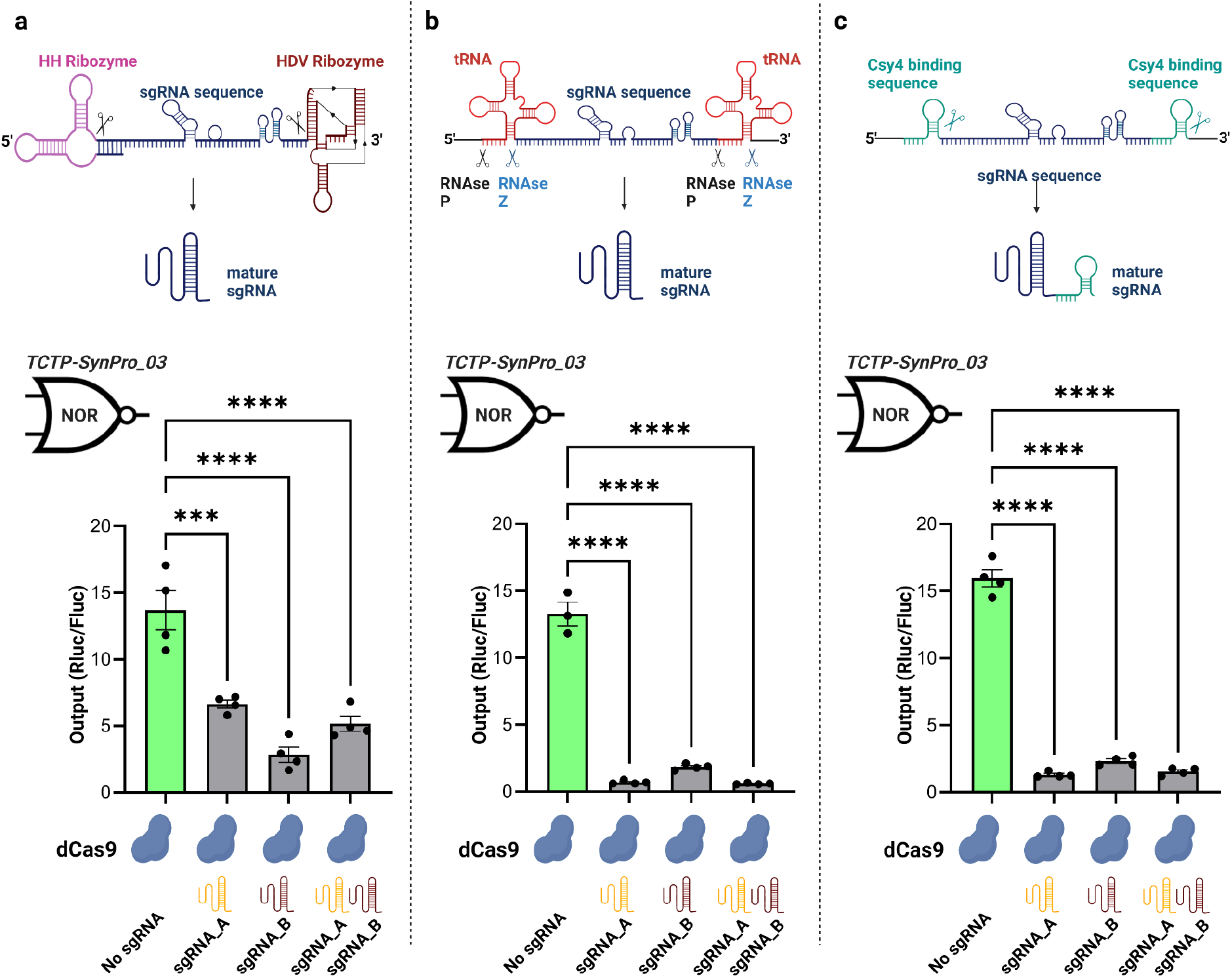
Comparison of sgRNA processing systems for Pol II-driven circuit input sgRNAs. sgRNAs were flanked by: **(a)** hammerhead (HH) and Hepatitis Delta Virus (HDV) ribozyme sequences, **(b)** tRNA (Gly) sequences, and **(c)** the Csy4 binding sequences. The whole sequence is placed immediately after the *CaMV 35S* promoter, and once processed the mature processed sgRNAs act as the input sequences targeting CRISPRi at *TCTP-SynPro_03* based programmable NOR gates. Output is relative luminescence (Rluc/Fluc). Asterisks (*** = p ≤ 0.0002 and **** = p ≤ 0.0001) indicate a significant difference compared to the No sgRNA using a one-way ANOVA and post hoc test (Tukey multiple comparison). Error bars represent standard error (n = 3 - 4).

The ribozyme-processed sgRNAs were able to produce NOR logic, however they were less effective in repressing the *TCTP-SynPro_03* integrator (Fig. 3a) than the *U6* produced sgRNAs (Fig. 2d), reducing circuit output to 20-50% of the No sgRNA control (Fig. 3a). The tRNA-based sgRNA processing system showed superior NOR gate repression, reducing circuit output to 4-15% compared to when No sgRNA was present (Fig. 3b). However, the tRNA sequence is known to contain binding sites for transcription factor IIIC (TF IIIC), which recruits TF IIIB and RNA Pol III^39^, and therefore there is a possibility of sgRNA transcription being driven by Pol III, regardless of the activation state of the upstream Pol II promoter. We tested whether tRNA-processed sgRNAs that lacked an upstream promoter could repress circuit output, revealing 53% repression in the absence of a Pol II promoter (Supplementary Fig. 3). This constitutes a potential problem for spatiotemporal control of gene expression, as the tRNA sequences flanking the sgRNA may potentially initiate Pol III transcription independently of any cell type-specific, conditional, or inducible Poll II promoters that may be used to drive input sgRNA expression and controlling circuit activity.

The performance of the Csy4-processed sgRNAs expressed from the *CaMV 35S* promoter was similar to the sgRNAs expressed from the *U6* promoter (Fig. 2d), causing repression of the *TCTP-SynPro_03* integrator to 8-15% compared to when no sgRNA was present (Fig. 3c). Thus, Csy4 provided equal or better performance than both the tRNA and ribozyme systems respectively, for processing input sgRNAs from Pol II promoters, and could be suitable for spatiotemporally-regulated expression of sgRNAs to integrate with programmable logic gates *in planta*. Crucially, effective processing of a sgRNA when it is expressed under a Pol II promoter also enables the transcriptional output of a NOR gate integrator to be yet another sgRNA, rather than simply a reporter gene. Such a sgRNA can in turn act as an input sgRNA targeting a different NOR gate integrator, enabling the connection and layering of multiple NOR gates to create more complex circuit logic. Thus, each NOR gate can act as a module, and input/output sgRNA signals can be sent and received between modules.

### Creating multi-layered complex logic gates

Creating more complex Boolean logic operations requires the connection of multiple logic gates in series. To produce the required logic expression, the output of the Layer 1 gate must display a clear separation between ON and OFF states, otherwise low levels of transcription of the output sgRNA might result in an incorrect logic state of the Layer 2 gate. In addition, the repression of the integrator must be dynamically reversible so that downstream gates are de-repressed when input sgRNAs target repression of the promoter of the upstream gate.

To demonstrate modularity and layering in our system, we aimed to create multi-layered logic gates. To ensure that integrator Layer 2 output was similar to the integrator Layer 1, we created additional synthetic *TCTP* promoters that have similar levels of activity in the absence of targeting sgRNAs by inserting new sgRNA target sequences flanking the TATA box (Supplementary Fig. 4).

We constructed an OR gate by connecting two different NOR gates together, with the first NOR-1 gate feeding into NOR-2 gate that controls the expression of *Renilla* luciferase (Fig. 4a). For this purpose, the output of our programmable NOR gate (Fig. 3c) was replaced with a new sgRNA (“sgRNA-C”), which targets the NOR-2 (*TCTP-SynPro_01)* integrator driving the expression of *Renilla* luciferase. We assembled all these components on a single plasmid and tested with all four possible combinations of the two input sgRNAs, demonstrating the correct OR logical functionality in plant cells (Fig. 4a and Supplementary Fig. 5a). In the absence of any input sgRNAs, sgRNA-C is expressed from integrator Layer 1 (*TCTP-SynPro_03*) to repress integrator Layer 2 (*TCTP-SynPro_01*), resulting in ∼74% repression compared to the ON state when input sgRNA-A and -B are present. However, we previously observed up to 92% repression with the NOR gate (Fig. 3c). Therefore, we sought to determine whether this is due to a lower efficacy of sgRNA-C or the strength of the *TCTP-SynPro_03* integrator driving sgRNA-C. We expressed sgRNA-C from the stronger *CaMV 35S* promoter to target the *TCTP-SynPro_01* integrator and found that the repression strength of sgRNA-C was significantly improved, resulting in 93% repression compared to the No input sgRNA (Supplementary Fig. 6). Thus, the amount of sgRNA produced from a Pol II promoter is an important design factor for producing effective CRISPRi in plant cells, and therefore balancing sgRNA production at different circuit layers through the choice of promoters in a multi-layered logic gate is important. Nevertheless, successful OR logic was produced with the *TCTP* driven 2-layer NOR design.

**Figure 4:**
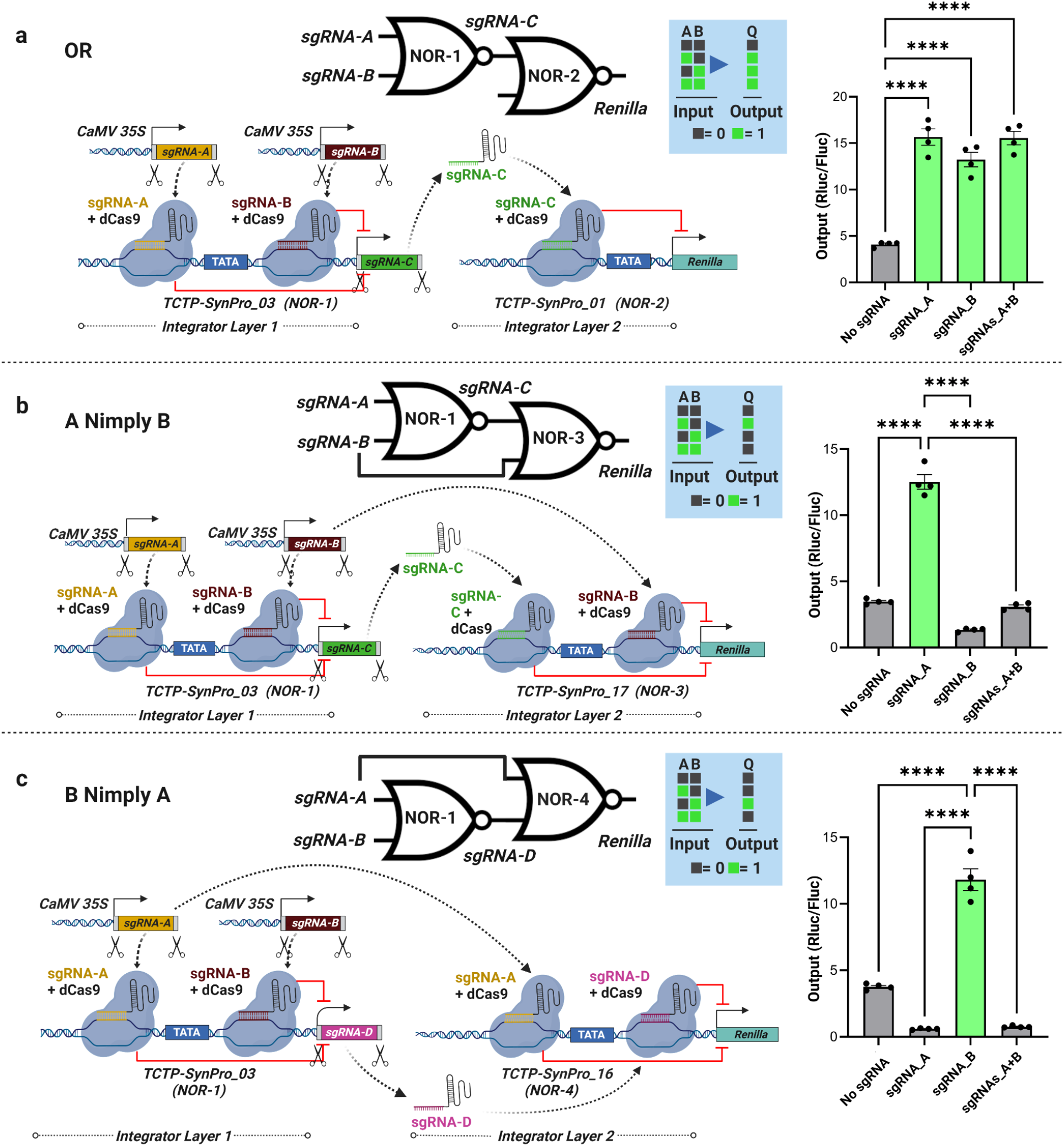
Programmable multi-layered circuits. Logic symbol, truth table, circuit design schematic, and measured output of the constructed circuit under different input signal conditions, for CRISPRi-based **(a)** OR, **(b)** A Nimply B, and **(c)** B Nimply A logic gates. Scissors represent Csy4 used for processing sgRNAs from the *CaMV 35S* promoters and Integrators of Layer 1. Output is relative luminescence (Rluc/Fluc) at 36 hpt. Asterisks (**** = p ≤ 0.0001) indicate a significant difference compared to the ON state using one-way ANOVA and post hoc test (Tukey multiple comparison test). Error bars represent standard error, n = 4.

To create an A Nimply B logic gate, we used the same Layer 1 integrator driving the expression of sgRNA-C from the OR gate. However, for Layer 2 (NOR-3) we designed a new integrator (*TCTP-SynPro_17*; Supplementary Fig. 3) to drive the expression of *Renilla* luciferase (Fig. 4b). This new integrator contains binding sites for both sgRNA-B and sgRNA-C, and therefore can be switched off in the presence of either of these two sgRNAs. This means that the A Nimply B gate will only produce Rluc output when sgRNA-A is provided alone, as sgRNA-B would act to repress integrator Layer 2, regardless of the presence of sgRNA-A, and override its stimulatory activity on the circuit. To test this new design of layering the NOR gates, we assembled all the circuit components on a single plasmid. Testing in the protoplast system demonstrated that the correct A Nimply B logic function was achieved (Fig. 4b and Supplementary Fig. 5b).

For the construction of a B Nimply A logic gate, in integrator Layer 1 we used *TCTP-SynpPro_03* to drive the expression of sgRNA-D, whereas for integrator Layer 2 (NOR-4) we designed a new integrator (*TCTP-SynPro_16*; Supplementary Fig. 3) containing binding sites for sgRNA-A and sgRNA-D, to drive the expression of *Renilla* luciferase (Fig. 4c). By testing this new layering combination of the NOR gates in protoplasts (all components assembled on a single plasmid), we achieved the correct B Nimply A logic functionality (Fig. 4c and Supplementary Fig. 5c). Thus, our results demonstrate that our programmable NOR gates can be layered in different ways to create complex circuits.

To determine whether we can combine more than two NOR gates together to produce logical operations, we created an AND gate by linking three different NOR gates together. The initial design showed suboptimal performance when input sgRNA-B was expressed (Supplementary Fig. 7a). We hypothesised that this difference might be because *TCTP-SynPro_18* (NOR-5) is not as strong as *TCTP-SynPro_19* (NOR-6; Supplementary Fig. 3), and therefore in the presence of input sgRNA-B the amount of sgRNA-C produced from NOR-5 is insufficient to effectively repress the integrator of Layer 2 (*TCTP-SynPro_01*) that drives *Renilla* luciferase expression. To resolve this, we created Versions 2 and 3 of the AND gate (Supplementary Figs. 6b and c). In both versions we used integrators of similar strengths within the Layer 1 (*TCTP-SynPro_16* [NOR-4] and *TCTP-SynPro_17* [NOR-3]), however we chose weaker integrators for the Layer 2 to see whether these could be repressed more effectively (*TCTP-SynPro_04* for Version 2 and *TCTP-SynPro_06* for Version 3; Supplementary Fig. 3). While the performance of input sgRNA-B in Versions 1-3 was similar at 24 hpt, Version 2 and Version 3 AND gates performed worse at 48 hpt, with no significant difference observed between input B and input A+B states (Supplementary Figs. 7b and c).

As mentioned previously, we used a single plasmid to encode all components of our dCas9-based logic circuits to ensure all components are present in each transfected cell. Because the *CaMV 35S* promoter that expresses the input sgRNA-B is sufficient to repress the *TCTP-*based integrators in previously built logic gates (Figs. 2-4), we hypothesised that the architecture of the components within the plasmid may be interfering with the performance of the constructs. To address this, we linearized the Version 2 AND gate by digesting with the restriction enzyme NotI to physically separate the Layer 1 integrators (Supplementary Fig. 8). Interestingly, circuit output repression induced by the input sgRNA-B was significantly improved, with repression similar to the input sgRNA-A (Fig. 5). This result suggests that plasmid architecture is also an important consideration for construction of effective logic gates, and overcoming this can achieve AND gate logic with our CRISPRi gene circuit designs.

**Figure 5:**
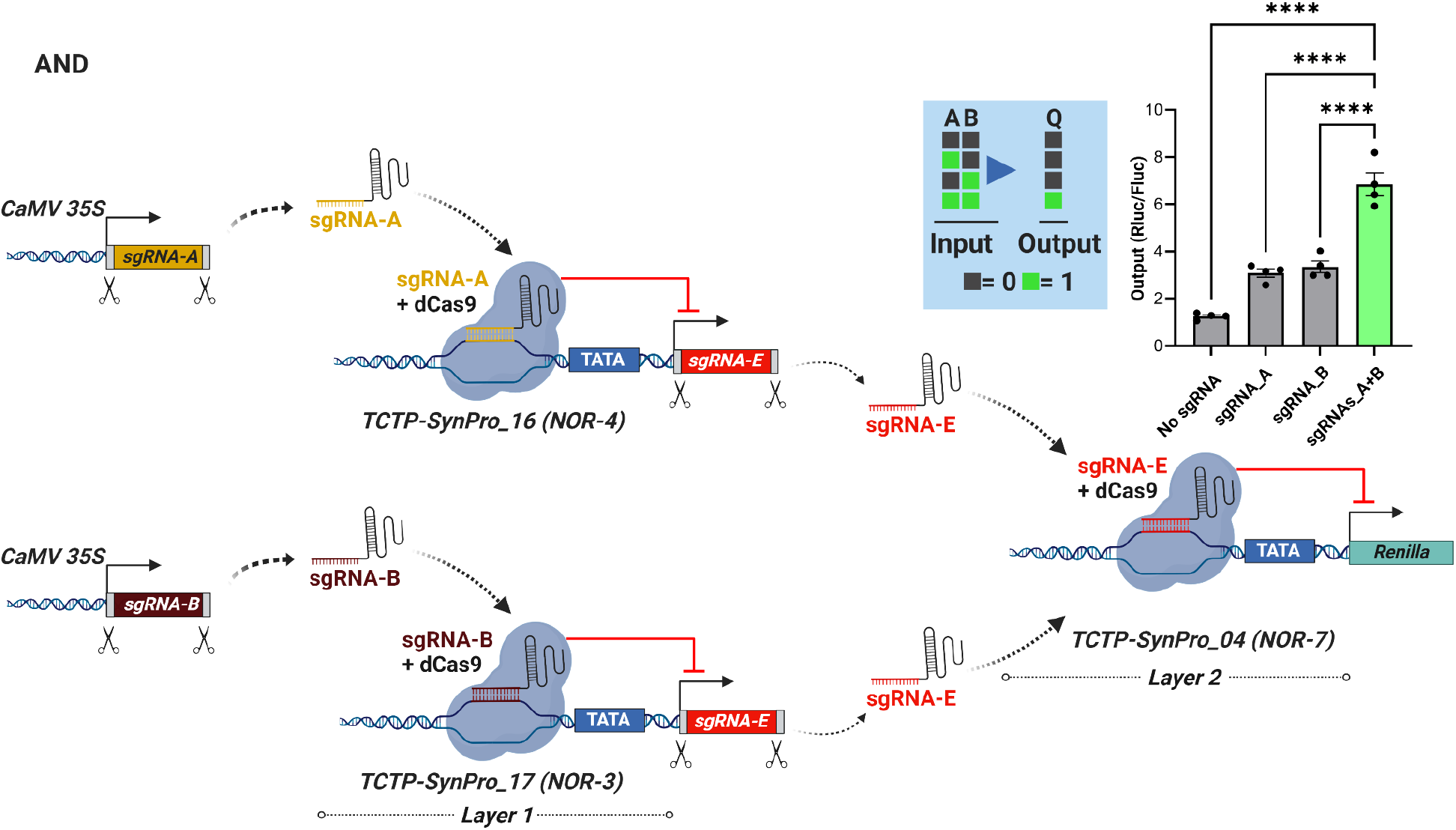
Implementation of a programmable AND gate. Logic symbol, truth table and schematic of circuit design for the implementation of the Version 2 of AND gate with a linearized input B plasmid. Processing of the sgRNAs from *CaMV 35S* promoters and Integrators via Csy4 is indicated by scissors. Output is the relative luciferase activity (Rluc/Fluc). Asterisks (**** = p ≤ 0.0001) indicate a significant difference compared to the ON state using one-way ANOVA and post hoc test (Tukey multiple comparison test). Error bars = standard error and n = 4.

## Discussion

In this work, for the first time in plants, we have demonstrated the successful implementation of programmable CRISPRi-based gene circuits. Using a high-throughput protoplast transfection protocol, we engineered the *TCTP* promoter to form a CRISPRi-based integrator, and successfully extended this to the widely used *CaMV 35S* promoter, suggesting that our designs can be applied to other promoters of choice. We demonstrated engineered promoter strength tunability, created new sets of variable-strength synthetic promoters for use as logic gate integrators, constructed NOR gates and demonstrated their correct activity in plant cells, and identified an effective sgRNA processing system to produce functional mature sgRNAs expressed from Pol II promoters to enable integration of our logic gates with endogenous input promoters and connected multiple logic gates modules together to produce additional Boolean logic gates. By connecting different NOR gates together, we created multi-layered OR, A Nimply B, B Nimply A, and AND logic gates to demonstrate modularity of this system.

Our CRISPRi circuit design has significant advantages over past systems that use different TF DNA binding domains to specify different binding events in a circuit^14,15,26^. Essentially, the circuit logic is programmed simply by changing the input sgRNAs and target sequences and linking the compact integrator modules (∼650 bp each) together, providing a very high level of flexibility. This has substantial advantages compared to circuit designs that require different recombinases due to the compactness of the system (recombinases can be ∼1.5 - 2kb in length) ^16^ or complex synthesis and delivery of many different DNA binding domains^15,26^, with potentially challenging and limited reprogrammability. Importantly, the integrator components of our circuits do not require the use of repurposed plant TFs, providing greater orthogonality from endogenous plant regulatory processes that may vary between cell types and conditions. Furthermore, our system avoids the potential off-target regulatory effects of engineered DNA binding domains linked to strong transcriptional activators or repressors that can influence transcription at a distance, which could increase the number of off-target transcription sites throughout the genome that elicit unintended regulatory effects^40^. Lastly, sgRNAs in the integrator are easily re-programmed and therefore can be replaced with different sgRNAs to reduce off-target events more easily as compared to TF or recombinase based circuits.

The sets of engineered *TCTP* and *CaMV 35S* promoters of different strengths reported here will be a useful resource for the plant synthetic biology community. The difference in strength of these modified promoters is likely due to the sgRNA binding sequences that replaced the native sequences flanking the TATA box. This may result in the deletion of natural TF binding sites, introduction of new binding sites for native TFs^41^, affect nucleosome occupancy^42^ or affect the binding strength of general transcription factors such as the TATA Binding Proteins 1 and 2 (TBP1 and TBP2) in *Arabidopsis*^43,44^. Our results are similar to the findings of Gander *et al*.^20^, where the activity of the upstream *GDP* promoter in *S. cerevisiae* was affected by replacing the native sequence of the core promoter with sgRNA binding sites.

The Csy4-processed sgRNAs from Pol II promoters were efficient in achieving repression to create a NOR gate, and will enable spatiotemporal control of sgRNA expression. However, efficient NOR integrator repression will likely depend on the quantity of sgRNAs produced from cell type-specific and inducible promoters. Therefore, to deploy CRISPRi-based logic gates for spatiotemporal control of gene expression *in vivo* it will be necessary to test multiple inducible and cell type-specific promoters for processing of sgRNAs and determine their performance in achieving single and multi-layered logic functions.

We have constructed 1, and 2 layer NOR logic gates to achieve complex Boolean logical operations, suggesting that our designed multi-layered circuits perform in a predictable manner without any signal degradation. However, we observed suboptimal performance of the input B in different versions of the two layered AND gate, resolved by linearizing the plasmid DNA to physically separate the NOR-4 and NOR-3 components, suggesting some interference from upstream DNA sequences. The inclusion of more effective insulators or independent insertion of NOR gates in transgenic plants would likely solve this. Therefore, similar to TF based circuits which have also shown unexpected outcomes during the testing and debugging phase^45^, optimization of CRISPRi based logic gates is necessary prior to implementation *in vivo*.

Overall, we have developed the first CRISPRi-based genetic logic gates in plants. CRISPRi was shown to be effective in repressing strong and moderate promoters in plant protoplasts, and therefore provides a new platform for the construction of compact, modular, and reversible gene circuits to achieve Boolean logic operations and complex programmable control of gene expression in plant cells. The toolkit of engineered promoters and gene circuits presented in this work will further advance plant synthetic biology and enable the implementation of more sophisticated and deliberate spatiotemporal control of plant gene regulatory pathways and cellular functions.

## Materials and Methods

### Plasmid construction

All the plasmids generated in this study were cloned using restriction and Gibson assembly based cloning^46^. All the genetic parts (promoters, CDS, terminators) were ordered from Integrated DNA Technologies (IDT) as gBlock synthetic gene fragment and cloned into the commercially available *pBluescript SK*(+) plasmid. Full sequences of plasmids used in this study are available on [Zenodo](https://doi.org/10.5281/zenodo.6783186).

### Plasmid sequencing

All plasmids in this manuscript were sequenced prior to testing. Whole plasmid sequencing was performed using Illumina Miseq. For this purpose, 100ng of plasmid DNA was tagmented using Tn5 enzyme. PCR was performed (10 cycles) to amplify the tagmented DNA with sequencing adapters using 2x MyTaq (BIO-25041; Bioline). The PCR products were cleaned using Serapure beads and pooled together prior to sequencing. Once sequenced, Unicycler^47^ and Bowtie^48^ were used for the *De novo* assembly of reads into a plasmid sequence or alignment to a reference plasmid sequence, respectively.

### Protoplast isolation and transfections in a 96-well plate

For protoplast isolation, *Arabidopsis* plants of the Columbia (Col-0) strain were grown on soil at 22°C in long day conditions (16 hours light / 8 hours dark). The mesophyll protoplasts were isolated from the leaves of 3 - 4 weeks old plants using the “Tape-Arabidopsis Sandwich” protocol^49^. For transfections in a 96-well plate, 5μg of the plasmid DNA (in 5μl) was added to each well followed by the addition of 10,000 cells (in 50μl). Using a multi-channel pipette, plasmid DNA and protoplasts were mixed with 55μl of 40% polyethylene glycol (PEG) solution and incubated for 15 minutes at room temperature. Transfected protoplasts were kept in continuous light at 25 °C for the desired period of time (16-48 hours). For each construct, four replicates were used and plasmid DNA was added in a randomized manner to avoid any positional effect on the plate. A detailed step-by-step protoplast transfection protocol has been submitted to the protocols.io repository. The Dual-Luciferase Reporter Assay System (E1960; Promega) was used for both cell lysis and measuring luciferase activity as per the manufacturer’s instructions using a POLARstar OPTIMA (BMG LABTECH) plate reader.

### Figure preparation

GraphPad Prism 9 and Biorender were used to generate bar plots and figures, respectively.

## Acknowledgements

The plasmid-seq data was generated on instrumentation supported by the Australian Cancer Research Foundation Centre for Advanced Cancer Genomics and Genomics WA.

## Author Contribution

M.A.K., B.K. and R.L. conceived the study, designed the experiments, and wrote the manuscript. M.A.K. with B.K., G.H., M.O., E.F., J.Y.Z., and B.J. conducted the experiments. M.A.K. designed the constructs and developed the 96-well protoplast transfection protocol. J.P., T.S. and C.P. conducted the plasmid sequencing. J.L, D.S. and I.S. provided assistance with the initial designs of constructs and designing experiments. All authors approved of and contributed to the final version of the manuscript.

## Supplementary Information

**Supplementary Figure 1:**
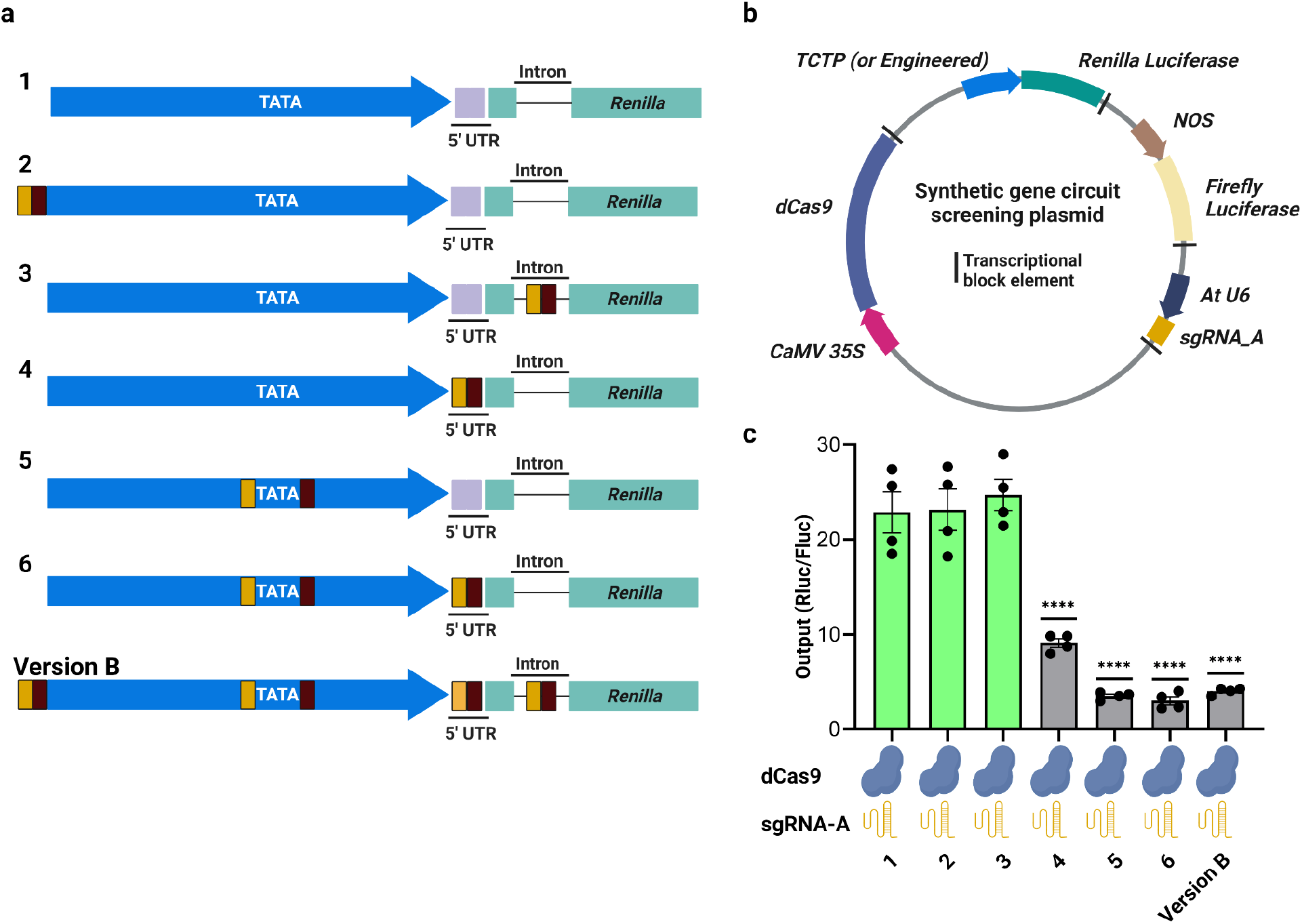
sgRNA target position within the *TCTP* promoter affects CRISPRi-based repression. **(a)** Designs of the wild-type or engineered *TCTP* promoters with sgRNA-A and sgRNA-B target sites (yellow and brown boxes, respectively) incorporated at different positions in the *TCTP* promoter and intron of the *Renilla* luciferase reporter. **(b)** Schematic representation of the screening plasmid (used for testing the promoters as per A) containing all the components required for testing CRISPRi in plant cells. **(c)** CRISPRi at different regions of the promoter compared to the presence (Version B) and absence of all binding sites (construct 1). Output is the relative luminescence ratio (Rluc/Fluc). Asterisks (**** = p ≤ 0.0001) indicate a significant difference compared to the control (construct 1) using one-way ANOVA and post hoc test (Tukey multiple comparison test). Error bars represent standard error, n = 4.

**Supplementary Figure 2:**
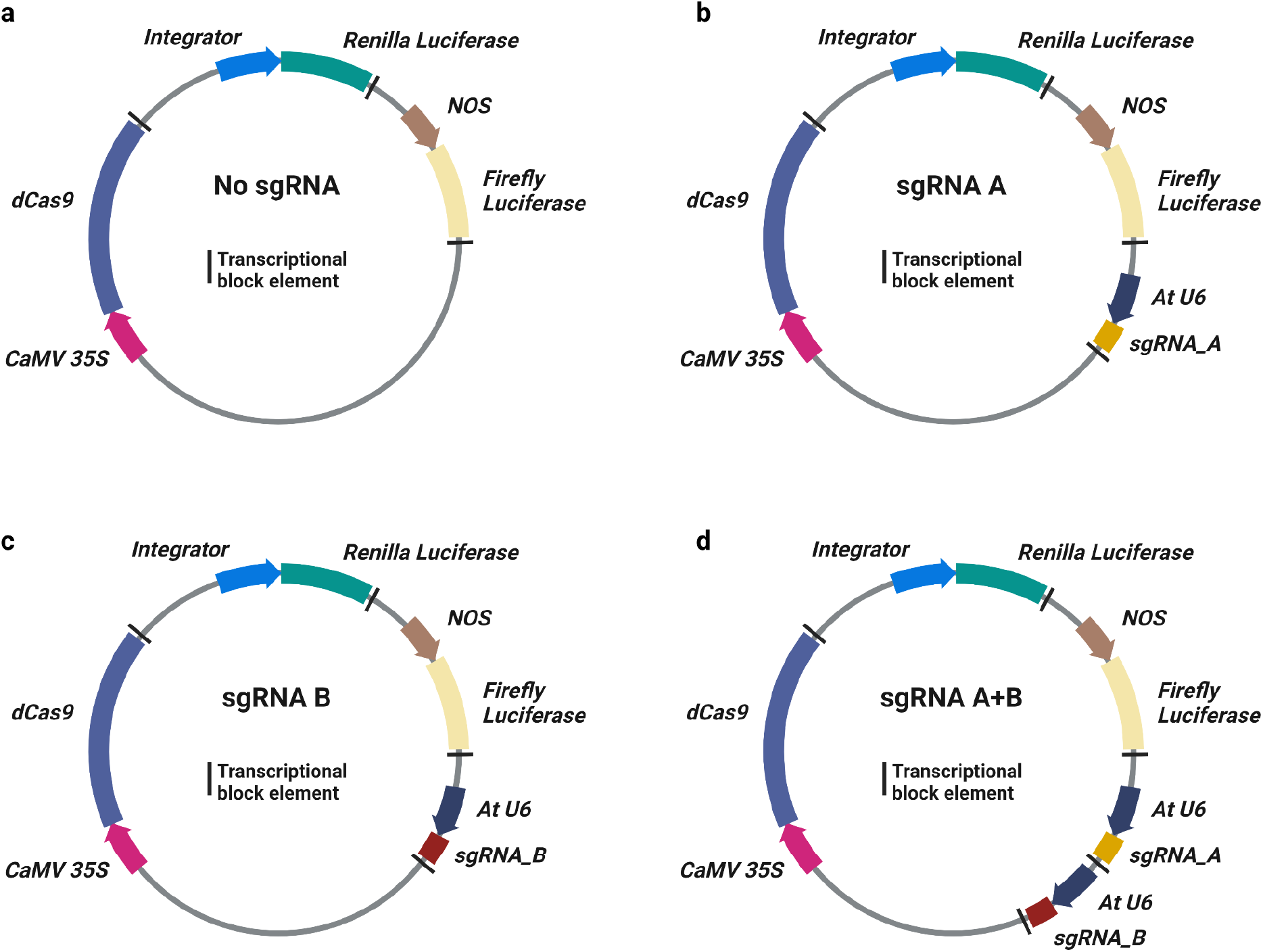
Screening plasmids used for testing NOR gates in plant cells. Schematic representation of the **(a)** No sgRNA, **(b)** input sgRNA-A, **(c)** input sgRNA-B, and **(d)** input sgRNAs-A+B used for testing NOR gates in plant cells. The integrator used for driving the expression of *Renilla* luciferase gene represents *TCTP-SynPro_03* or *CaMV 35S-SynPro_01*.

**Supplementary Figure 3:**
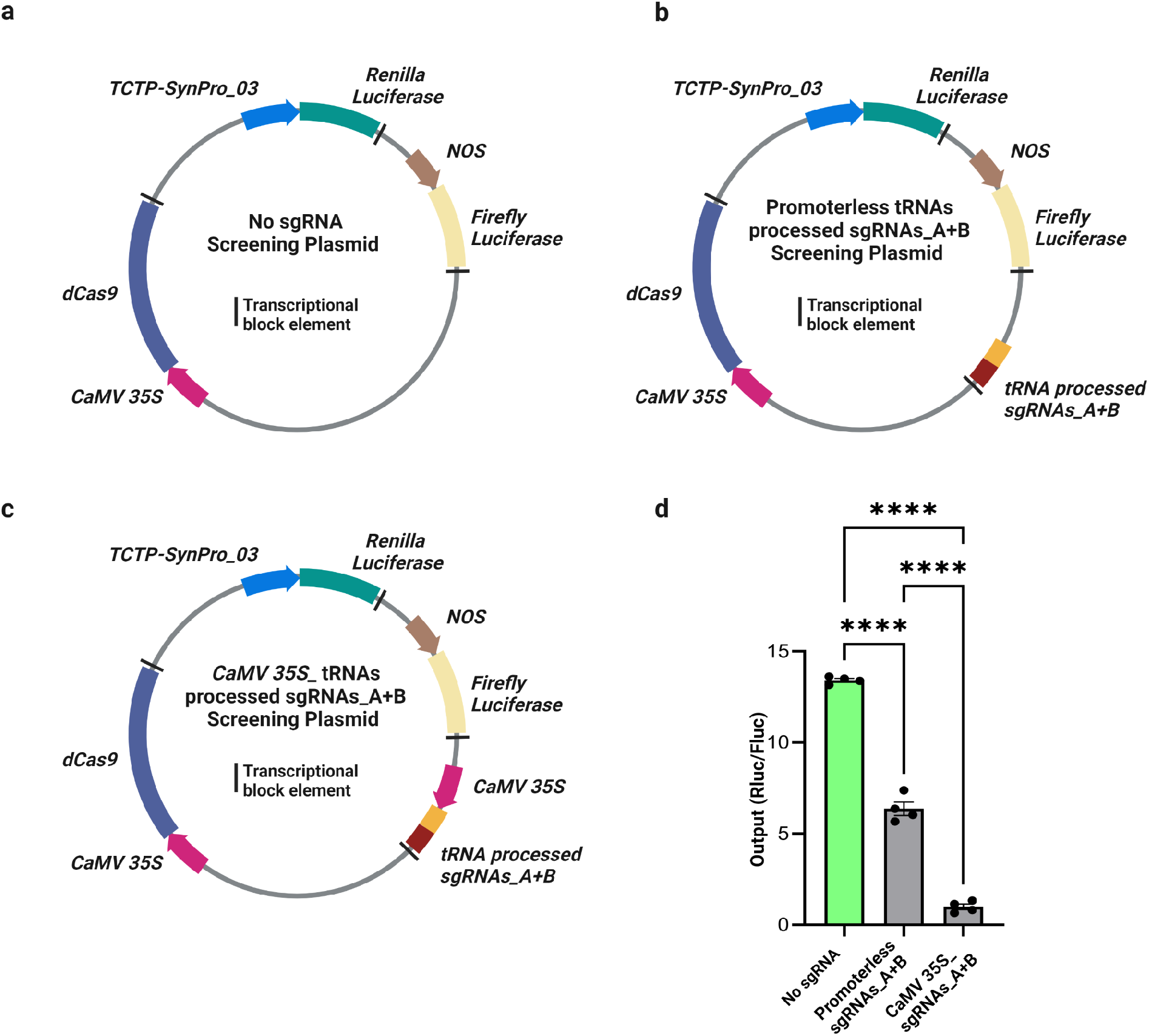
tRNA-processed sgRNAs inhibit *TCTP-SynPro_03* integrator activity in the absence of an upstream Pol II promoter. Schematic representation of screening plasmids used for **(a)** No sgRNA, **(b)** promoterless tRNAs processed sgRNAs_A+B, and **(c)** *CaMV 35S*_tRNAs processed sgRNAs_A+B. **(d)** Comparison of tRNA-processed sgRNAs in the presence or absence of an upstream *CaMV 35S* promoter driving expression. In the absence of the Pol II promoter the tRNA-processed sgRNAs reduce circuit output to 53% of the No sgRNA control. Output is the relative luciferase activity (Rluc/Fluc). Asterisks (**** = p ≤ 0.0001) indicate a significant difference using one-way ANOVA and post-hoc test (Tukey multiple comparison test). Error bars represent standard error, n = 4.

**Supplementary Figure 4:**
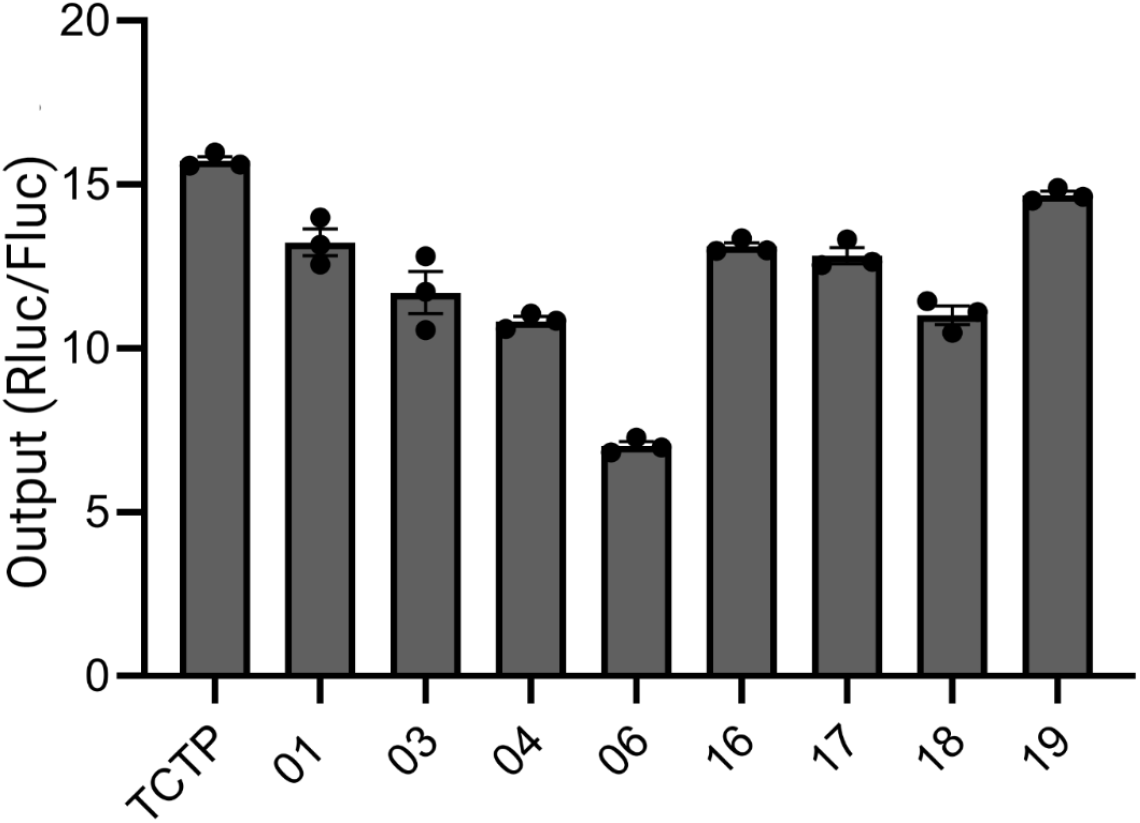
Activity of engineered *TCTP* promoters used for the construction of multi-layered circuits. Bar plot comparing output (Rluc/Fluc) of the additional engineered *TCTP* promoters used for multi-layered circuits compared to the first library of synthetic TCTP promoters (n = 3).

**Supplementary Figure 5:**
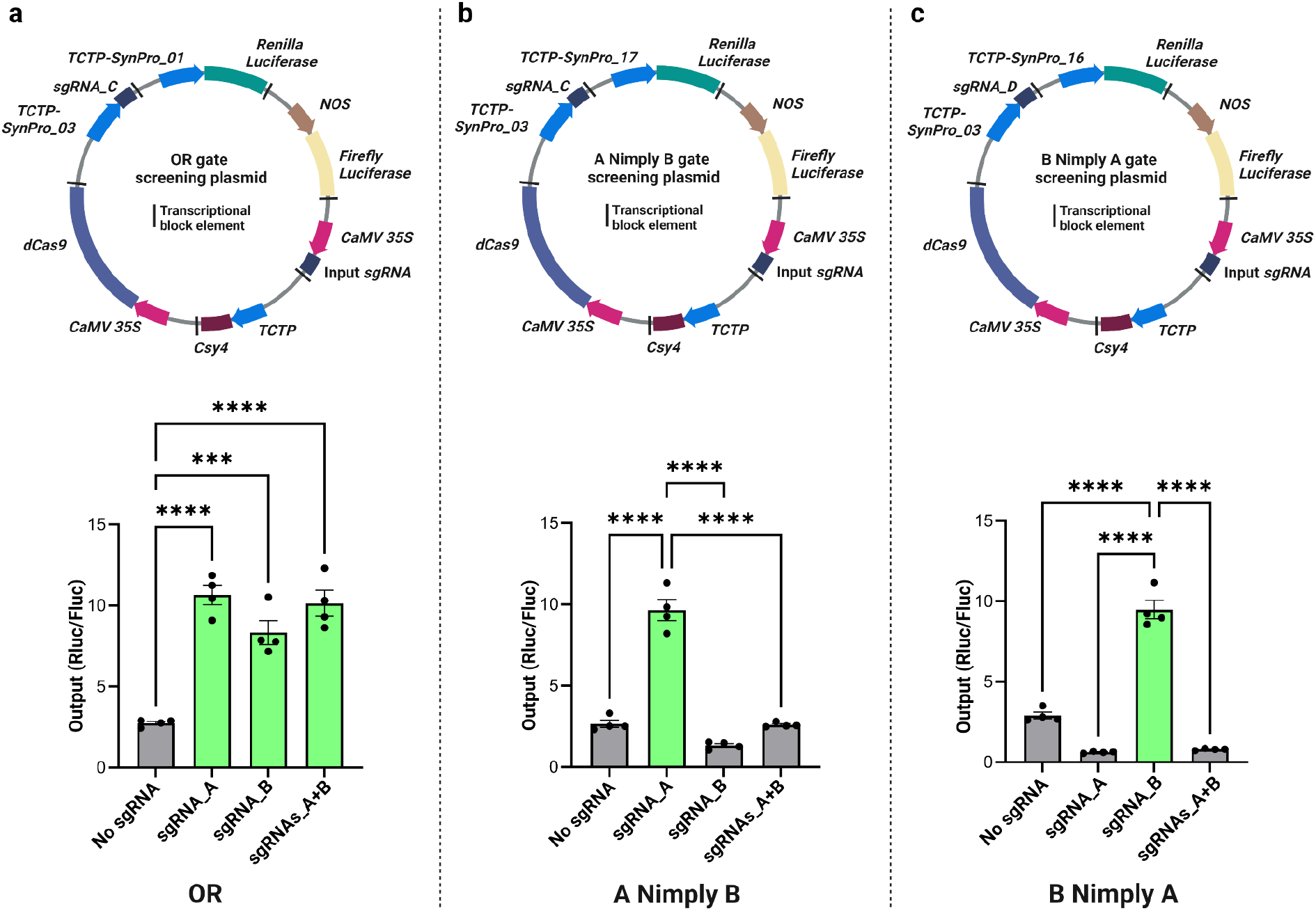
Performance of OR, A Nimply B, and B Nimply A multi-layered circuits at 24 hpt. Schematic representation of screening plasmids and performance of **(a)** OR, **(b)** A Nimply B, and **(c)** B Nimply A gates in plant cells at 24 hpt. For each gate four different plasmids were constructed that contained all the components required for testing each with *CaMV 35S* processed Input sgRNA-A, -B, -A+B, or No sgRNA. Output represents the relative luminescence ratio (Rluc/Fluc). Asterisks (*** = P ≤ 0.001, **** = P ≤ 0.0001) indicate a significant difference compared to the ON state using one-way ANOVA and post hoc test (Tukey multiple comparison test). Error bars represent standard error, n = 4.

**Supplementary Figure 6:**
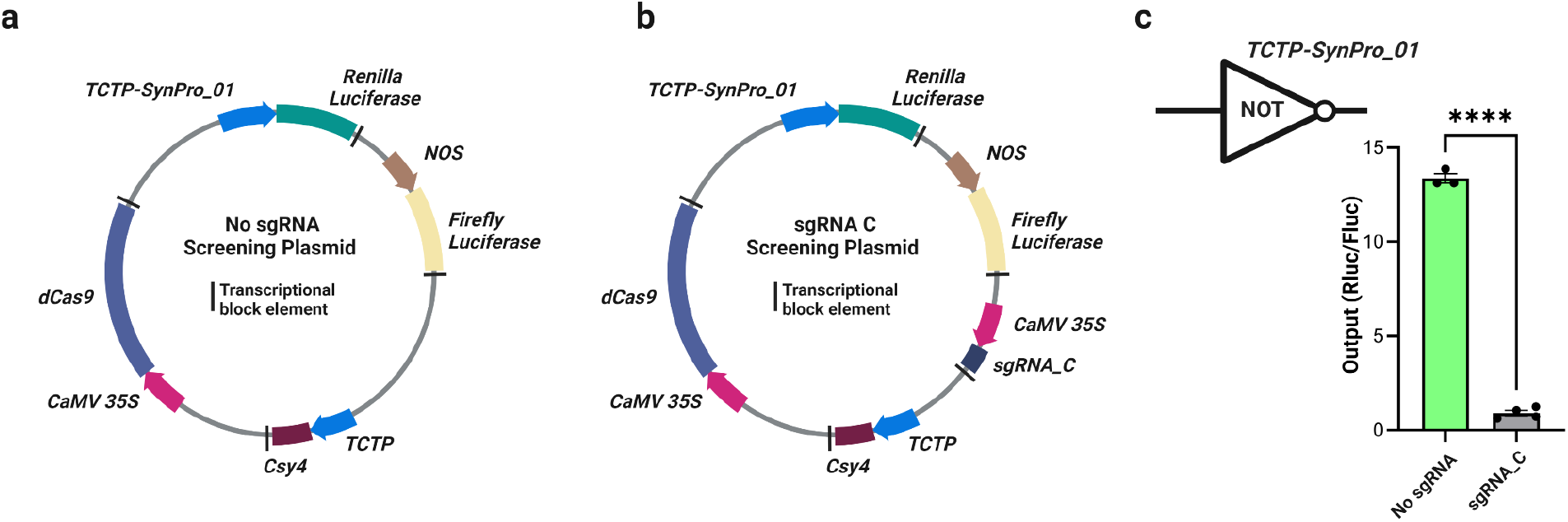
Promoter inhibition by sgRNA-C when it is expressed from a strong Pol II promoter. Constructs used for **(a)** No sgRNA and **(b)** input sgRNA-C for testing in plant cells at 24 hpt. The sgRNA-C is expressed under the *CaMV 35S* promoter and processed using the Csy4 endonuclease system, resulting in 93% repression **(c)**. Output is the relative luminescence (Rluc/Fluc). Asterisks (**** = p ≤ 0.0001) indicate significant difference compared to no input sgRNA using two-tailed student’s t-tests. Error bar, represents standard error, n = 3-4.

**Supplementary Figure 7:**
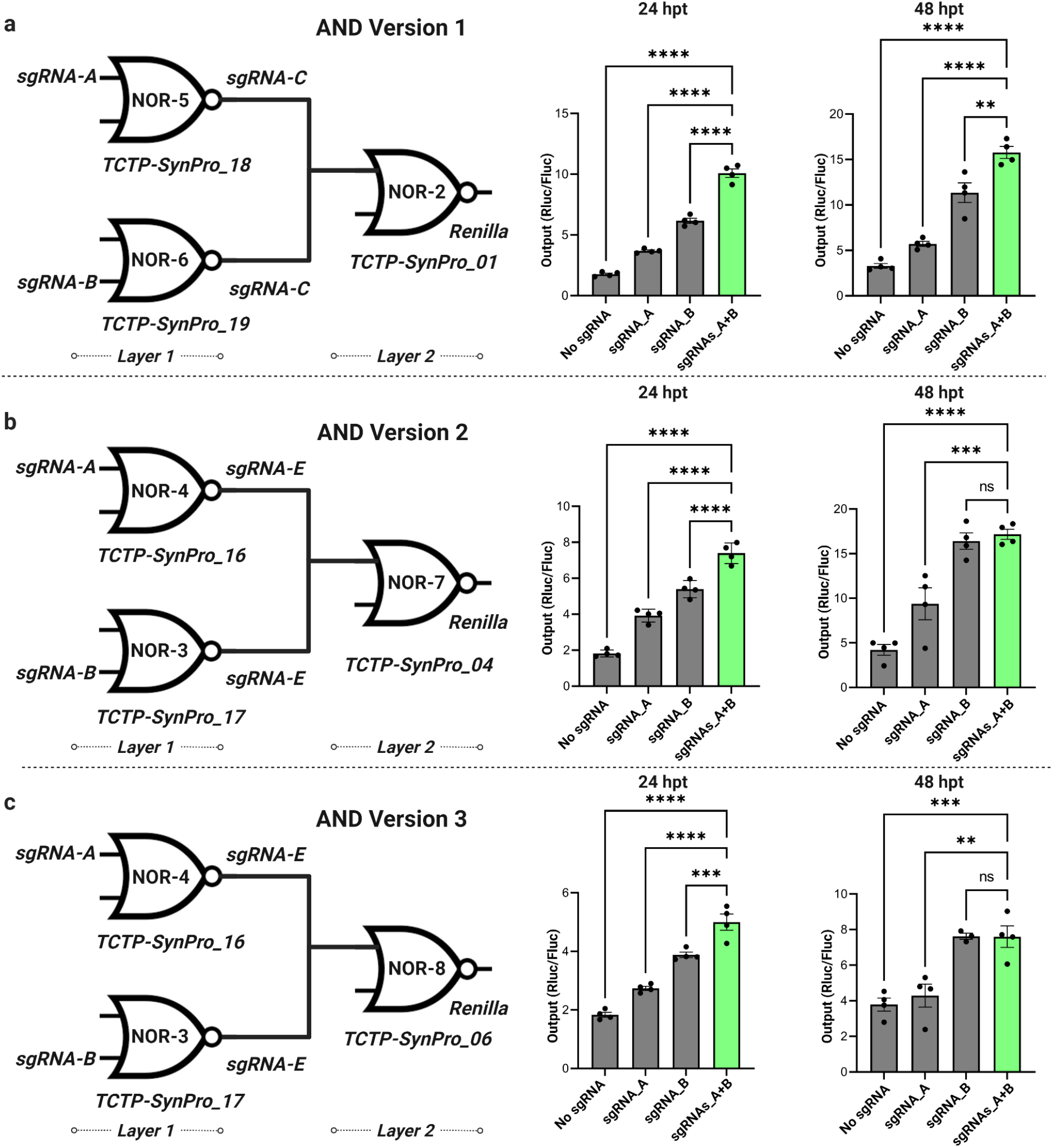
Performance of different versions of AND gates in plant cells. Output is the relative luciferase activity (Rluc/Fluc). Asterisks (** = P ≤ 0.01, *** = P ≤ 0.001, **** = P ≤ 0.0001) indicate a significant difference compared to the ON state using one-way ANOVA and post hoc test (Tukey multiple comparison test). Error bars represent standard error, n = 4.

**Supplementary Figure 8:**
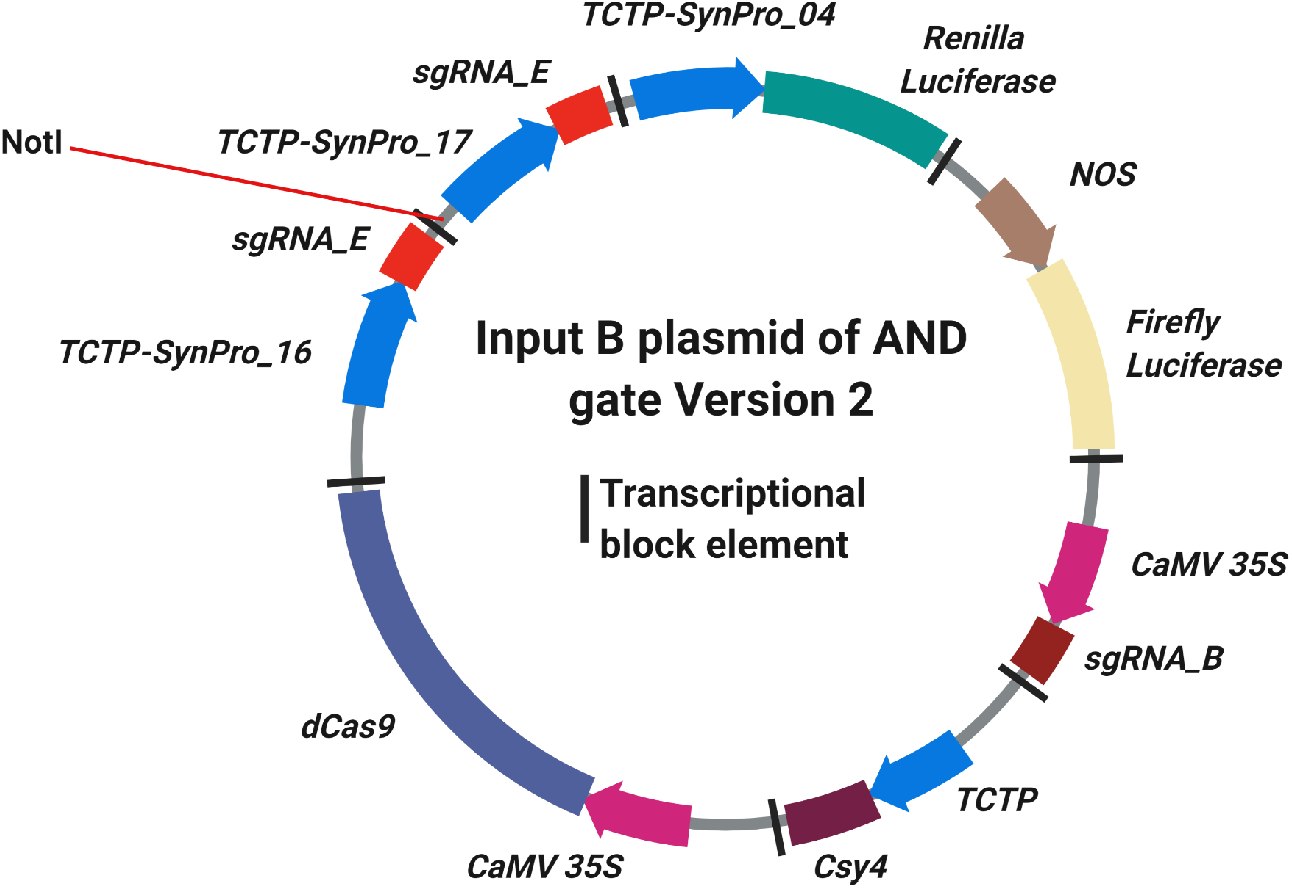
Schematic representation of the Input B plasmid of the AND gate version 2.

**Supplementary Table 1:**
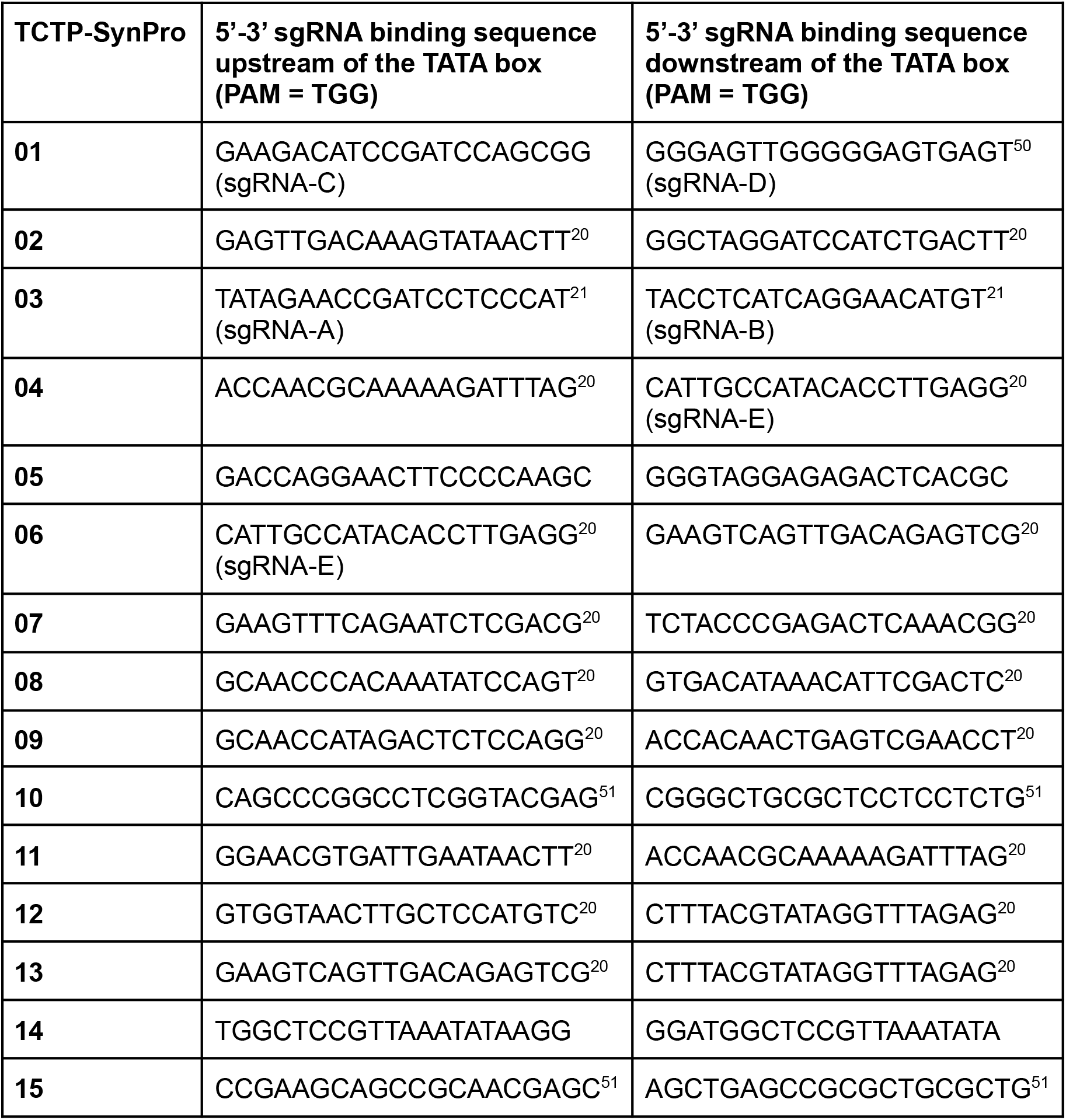
sgRNA binding sequences used for creating TCTP-based synthetic promoters

**Supplementary Table 2:**
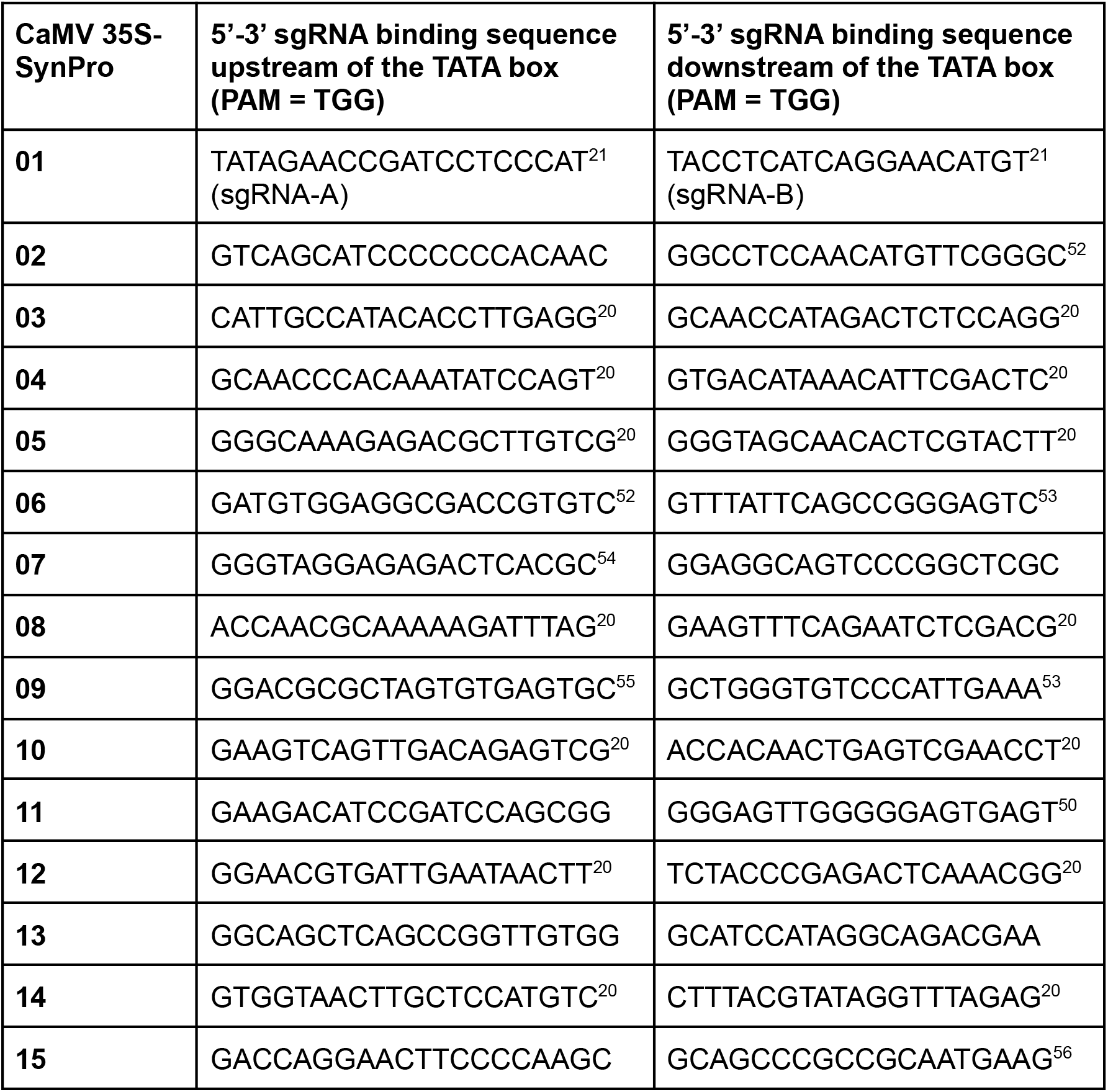
sgRNA binding sequences used for creating CaMV 35S-based synthetic promoters

## References

1. Thompson, A. J. et al. Ectopic expression of a tomato 9-cis-epoxycarotenoid dioxygenase gene causes over-production of abscisic acid. Plant J. 23, 363–374 (2000).

2. Iuchi, S. et al. Regulation of drought tolerance by gene manipulation of 9-cis-epoxycarotenoid dioxygenase, a key enzyme in abscisic acid biosynthesis in Arabidopsis. Plant J. 27, 325–333 (2001).

3. Feeney, M., Frigerio, L., Cui, Y. & Menassa, R. Following vegetative to embryonic cellular changes in leaves of Arabidopsis overexpressing LEAFY COTYLEDON2. Plant Physiol. 162, 1881–1896 (2013).

4. Vanhercke, T. et al. Step changes in leaf oil accumulation via iterative metabolic engineering. Metab. Eng. 39, 237–246 (2017).

5. He, R. et al. Overexpression of 9-cis-Epoxycarotenoid Dioxygenase Cisgene in Grapevine Increases Drought Tolerance and Results in Pleiotropic Effects. Front. Plant Sci. 9, 970 (2018).

6. Brophy, J. A. N. & Voigt, C. A. Principles of genetic circuit design. Nat. Methods 11, 508–520 (2014).

7. Kassaw, T. K., Donayre-Torres, A. J., Antunes, M. S., Morey, K. J. & Medford, J. I. Engineering synthetic regulatory circuits in plants. Plant Sci. (2018) doi:10.1016/j.plantsci.2018.04.005.

8. Andres, J., Blomeier, T. & Zurbriggen, M. D. Synthetic Switches and Regulatory Circuits in Plants. Plant Physiol. 179, 862–884 (2019).

9. de Lange, O., Klavins, E. & Nemhauser, J. Synthetic genetic circuits in crop plants. Curr. Opin. Biotechnol. 49, 16–22 (2018).

10. Xia, P.-F., Ling, H., Foo, J. L. & Chang, M. W. Synthetic genetic circuits for programmable biological functionalities. Biotechnol. Adv. 37, 107393 (2019).

11. Verbič, A., Praznik, A. & Jerala, R. A guide to the design of synthetic gene networks in mammalian cells. FEBS J. 288, 5265–5288 (2021).

12. Chen, Y. et al. Genetic circuit design automation for yeast. Nature Microbiology 5, 1349–1360 (2020).

13. Vazquez-Vilar, M. et al. GB3.0: a platform for plant bio-design that connects functional DNA elements with associated biological data. Nucleic Acids Res. 45, 2196–2209 (2017).

14. Schaumberg, K. A. et al. Quantitative characterization of genetic parts and circuits for plant synthetic biology. Nat. Methods 13, 94 (2015).

15. Brophy, J. A. N., Magallon, K. J., Kniazev, K. & Dinneny, J. R. Synthetic genetic circuits enable reprogramming of plant roots. bioRxiv 2022.02.02.478917 (2022) doi:10.1101/2022.02.02.478917.

16. Lloyd, J. P. B. et al. Synthetic memory circuits for programmable cell reconfiguration in plants. bioRxiv 2022.02.11.480167 (2022) doi:10.1101/2022.02.11.480167.

17. Belcher, M. S. et al. Design of orthogonal regulatory systems for modulating gene expression in plants. Nat. Chem. Biol. 16, 857–865 (2020).

18. Bernabé-Orts, J. M. et al. A memory switch for plant synthetic biology based on the phage ϕC31 integration system. Nucleic Acids Res. 48, 3379–3394 (2020).

19. Schürholz, A.-K. et al. A Comprehensive Toolkit for Inducible, Cell Type-Specific Gene Expression in Arabidopsis. Plant Physiol. 178, 40–53 (2018).

20. Gander, M. W., Vrana, J. D., Voje, W. E., Carothers, J. M. & Klavins, E. Digital logic circuits in yeast with CRISPR-dCas9 NOR gates. Nat. Commun. 8, 15459 (2017).

21. Kiani, S. et al. CRISPR transcriptional repression devices and layered circuits in mammalian cells. Nat. Methods 11, 723–726 (2014).

22. Yeo, N. C. et al. An enhanced CRISPR repressor for targeted mammalian gene regulation. Nat. Methods 15, 611–616 (2018).

23. Nielsen, A. A. K. & Voigt, C. A. Multi-input CRISPR/Cas genetic circuits that interface host regulatory networks. Mol. Syst. Biol. 10, 763 (2014).

24. Santos-Moreno, J., Tasiudi, E., Stelling, J. & Schaerli, Y. Multistable and dynamic CRISPRi-based synthetic circuits. Nat. Commun. 11, 2746 (2020).

25. Kim, H., Bojar, D. & Fussenegger, M. A CRISPR/Cas9-based central processing unit to program complex logic computation in human cells. Proc. Natl. Acad. Sci. U. S. A. 116, 7214–7219 (2019).

26. Gaber, R. et al. Designable DNA-binding domains enable construction of logic circuits in mammalian cells. Nat. Chem. Biol. 10, 203–208 (2014).

27. Lowder, L. G. et al. A CRISPR/Cas9 Toolbox for Multiplexed Plant Genome Editing and Transcriptional Regulation. Plant Physiol. 169, 971–985 (2015).

28. Piatek, A. et al. RNA-guided transcriptional regulation in planta via synthetic dCas9-based transcription factors. Plant Biotechnol. J. 13, 578–589 (2015).

29. Vazquez-Vilar, M. et al. A modular toolbox for gRNA-Cas9 genome engineering in plants based on the GoldenBraid standard. Plant Methods 12, 10 (2016).

30. Vazquez-Vilar, M. et al. The GB4.0 Platform, an All-In-One Tool for CRISPR/Cas-Based Multiplex Genome Engineering in Plants. Front. Plant Sci. 12, 689937 (2021).

31. Han, Y.-J., Kim, Y.-M., Hwang, O.-J. & Kim, J.-I. Characterization of a small constitutive promoter from Arabidopsis translationally controlled tumor protein (AtTCTP) gene for plant transformation. Plant Cell Rep. 34, 265–275 (2015).

32. Somssich, M. A short history of the CaMV 35S promoter. https://peerj.com/preprints/27096/ (2019) xdoi:10.7287/peerj.preprints.27096v3.

33. Vancanneyt, G., Schmidt, R., O’Connor-Sanchez, A., Willmitzer, L. & Rocha-Sosa, M. Construction of an intron-containing marker gene: splicing of the intron in transgenic plants and its use in monitoring early events in Agrobacterium-mediated plant transformation. Mol. Gen. Genet. 220, 245–250 (1990).

34. Gao, Y. & Zhao, Y. Self-processing of ribozyme-flanked RNAs into guide RNAs in vitro and in vivo for CRISPR-mediated genome editing. J. Integr. Plant Biol. 56, 343–349 (2014).

35. Cermak, T. et al. A multi-purpose toolkit to enable advanced genome engineering in plants. Plant Cell (2017) doi:10.1105/tpc.16.00922.

36. Xie, K., Minkenberg, B. & Yang, Y. Boosting CRISPR/Cas9 multiplex editing capability with the endogenous tRNA-processing system. Proc. Natl. Acad. Sci. U. S. A. 112, 3570–3575 (2015).

37. Haurwitz, R. E., Jinek, M., Wiedenheft, B., Zhou, K. & Doudna, J. A. Sequence- and structure-specific RNA processing by a CRISPR endonuclease. Science 329, 1355–1358 (2010).

38. Nissim, L., Perli, S. D., Fridkin, A., Perez-Pinera, P. & Lu, T. K. Multiplexed and programmable regulation of gene networks with an integrated RNA and CRISPR/Cas toolkit in human cells. Mol. Cell 54, 698–710 (2014).

39. Lodish, H. et al. Molecular Cell Biology. (W. H. Freeman, 2000).

40. Leben, K. et al. Binding of the transcription activator-like effector augments transcriptional regulation by another transcription factor. Nucleic Acids Res. 50, 6562–6574 (2022).

41. Tompa, M. et al. Assessing computational tools for the discovery of transcription factor binding sites. Nat. Biotechnol. 23, 137–144 (2005).

42. Jiang, C. & Pugh, B. F. Nucleosome positioning and gene regulation: advances through genomics. Nat. Rev. Genet. 10, 161–172 (2009).

43. Heard, D. J., Kiss, T. & Filipowicz, W. Both Arabidopsis TATA binding protein (TBP) isoforms are functionally identical in RNA polymerase II and III transcription in plant cells: evidence for gene-specific changes in DNA binding specificity of TBP. EMBO J. 12, 3519–3528 (1993).

44. Mukumoto, F., Hirose, S., Imaseki, H. & Yamazaki, K. DNA sequence requirement of a TATA element-binding protein from Arabidopsis for transcription in vitro. Plant Mol. Biol. 23, 995–1003 (1993).

45. Gorochowski, T. E. et al. Genetic circuit characterization and debugging using RNA-seq. Mol. Syst. Biol. 13, 952 (2017).

46. Gibson, D. G. et al. Enzymatic assembly of DNA molecules up to several hundred kilobases. Nat. Methods 6, 343–345 (2009).

47. Wick, R. R., Judd, L. M., Gorrie, C. L. & Holt, K. E. Unicycler: Resolving bacterial genome assemblies from short and long sequencing reads. PLoS Comput. Biol. 13, e1005595 (2017).

48. Langmead, B. & Salzberg, S. L. Fast gapped-read alignment with Bowtie 2. Nat. Methods 9, 357–359 (2012).

49. Wu, F.-H. et al. Tape-Arabidopsis Sandwich - a simpler Arabidopsis protoplast isolation method. Plant Methods 5, 16 (2009).

50. Cano-Rodriguez, D. et al. Writing of H3K4Me3 overcomes epigenetic silencing in a sustained but context-dependent manner. Nat. Commun. 7, 1–11 (2016).

51. Konermann, S. et al. Transcriptome Engineering with RNA-Targeting Type VI-D CRISPR Effectors. Cell 173, 665–676.e14 (2018).

52. Garcia-Bloj, B. et al. Waking up dormant tumor suppressor genes with zinc fingers, TALEs and the CRISPR/dCas9 system. Oncotarget 7, 60535–60554 (2016).

53. Shechner, D. M., Hacisuleyman, E., Younger, S. T. & Rinn, J. L. Multiplexable, locus-specific targeting of long RNAs with CRISPR-Display. Nat. Methods 12, 664–670 (2015).

54. Amabile, A. et al. Inheritable Silencing of Endogenous Genes by Hit-and-Run Targeted Epigenetic Editing. Cell 167, 219–232.e14 (2016).

55. Gilbert, L. A. et al. CRISPR-mediated modular RNA-guided regulation of transcription in eukaryotes. Cell 154, 442–451 (2013).

56. Maeder, M. L. et al. CRISPR RNA–guided activation of endogenous human genes. Nat. Methods 10, 977–979 (2013).

